# GenoM7GNet: An Efficient N^7^-methylguanosine Site Prediction Approach Based on a Nucleotide Language Model

**DOI:** 10.1101/2024.09.03.610976

**Authors:** Chuang Li, Heshi Wang, Yanhua Wen, Rui Yin, Xiangxiang Zeng, Keqin Li

**Affiliations:** School of Computer Science, Hunan University of Technology and Business, Hunan, China, 410205; Xiangjiang Laboratory, Hunan, China, 410205; Department of Health Outcomes and Biomedical Informatics, University of Florida, Gainsville, USA, FL 32608; College of Computer Science and Electronic Engineering, Hunan University, Hunan, China, 410082; College of Computer Science and Electronic Engineering, Hunan University, Changsha, China, 410082; Department of Computer Science, State University of New York, New Paltz, USA, NY 12561

**Keywords:** N^7^-methlguanosine, BERT, convolutional neural network, deep learning, biological sequence

## Abstract

N^7^-methylguanosine (m7G), one of the mainstream post-transcriptional RNA modifications, occupies an exceedingly significant place in medical treatments. However, classic approaches for identifying m7G sites are costly both in time and equipment. Meanwhile, the existing machine learning methods extract limited hidden information from RNA sequences, thus making it difficult to improve the accuracy. Therefore, we put forward to a deep learning network, called “GenoM7GNet,” for m7G site identification. This model utilizes a Bidirectional Encoder Representation from Transformers (BERT) and is pretrained on nucleotide sequences data to capture hidden patterns from RNA sequences for m7G site prediction. Moreover, through detailed comparative experiments with various deep learning models, we discovered that the one-dimensional convolutional neural network (CNN) exhibits outstanding performance in sequence feature learning and classification. The proposed GenoM7GNet model achieved 0.953 in accuracy, 0.932 in sensitivity, 0.976 in specificity, 0.907 in Matthews Correlation Coefficient and 0.984 in Area Under the receiver operating characteristic Curve on performance evaluation. Extensive experimental results further prove that our GenoM7GNet model markedly surpasses other state-of-the-art models in predicting m7G sites, exhibiting high computing performance.

## 1 Introduction

RNA modification refers to the chemical alteration of the nucleotide sequence of RNA molecules under specific conditions. It can enhance the capacity of RNA to convey diverse information, realizing more complex biological functions [1]. At present, among the identified RNA modifications, N^7^-methylguanosine (m7G) has become a rising star in the research field because of its significant epigenetic modifications [2], [3]. m7G refers to the methylation modification on the seventh nitrogen atom of the guanine in RNA molecules. Research has shown that m7G modification is ubiquitous in RNA molecules, including mRNA, ribosomal RNA (rRNA), and transfer RNA (tRNA), and it participates in numerous vital biological processes, including RNA processing, gene expression, and protein translation [4], [5], [6].

In previous studies, m7G modification has been shown to be highly correlated with a variety of diseases, such as Down syndrome, brain malformations, and microcephalic primordial dwarfism [4], [7], [8]. Recent research also indicates the involvement of m7G modification at various stages of cancer formation and progression [9], [10], [11]. In the study of mammals, Methyltransferase-like 1 (METTL1) is the most frequently researched m7G regulatory factor. Together with its co-factor WD Repeat Domain 4 (WDR4), it performed m7G modifications in mRNA, miRNA, and tRNA [12]. Usually, in cancer, abnormal expressions are found in m7G methyltransferases. They catalyze the m7G modification in miRNA or tRNA, affecting the expression of target genes, and thereby regulating the progression of tumors [13]. In summary, determining the distribution of m7G sites within RNA is fundamental for the in-depth study of related diseases and holds significant implications for medical treatment and research.

In traditional experimental methodologies, numerous techniques for identifying m7G sites have emerged due to the advancement of sequencing technology, including MeRIP-seq, miCLIP-seq, AlkAniline-seq, and m7G-quant-seq [6], [14], [15], [16]. However, these methods typically require extensive manipulations such as RNA enrichment, isolation, chemical reactions, and gel electrophoresis, rendering them time-consuming, laborious, and relatively costly. Moreover, the efficiency of traditional experimental methods is rather low for large-scale RNA sequencing. Consequently, there is an urgent need for an efficient prediction method to accurately identify m7G sites.

With increasingly available biological data and the rapid advancement of machine learning (ML), numerous researchers have applied new techniques to address problems in bioinformatics [17], [18], [19]. Chen *et al*. [20] pioneered the application of ML to identify m7G sites, proposing a predictor based on support vector machine (SVM) named iRNA-m7G. It utilized the integration approach of various features to enrich its information for prediction, including secondary structure component, nucleotide property and frequency, and pseudo nucleotide composition. Subsequently, Song et al. developed m7GFinder for the high-precision identification of m7G sites based on human genomic coordinates or DNA sequences [21]. Liu et al. presented m7GPredictor, which predicts m7G sites based on various feature extraction methods [22]. Other m7G site prediction tools include XG-m7G, m7G-IFL, m7G-DLSTM, THRONE, and m7G-autoBioSeqpy [23], [24], [25], [26], [27].

Despite the progress in the prediction of m7G sites with existing models, the latent spatial and temporal information of RNA sequences is underexplored, which could restrict the potential for further improvements in their prediction accuracy. Recently, there has been remarkable success in studies applying nucleotide language models to predict RNA-protein interactions [28] and DNA methylations [29]. Compared to previous feature extraction methods, nucleotide language models possess more potent capabilities for extracting features from nucleotide sequences, such as DNA and RNA sequences. We assumed that if we can capture a richer array of hidden information from RNA sequences, we could further enhance the accuracy and efficiency of m7G site prediction by integrating the nucleotide language model with deep learning.

In this paper, we proposed an efficient deep learning prediction model utilizing a nucleotide language model, named “GenoM7GNet.” GenoM7GNet primarily comprises two parts: a pre-trained BERT [30] model and a CNN model. In the pre-trained BERT model part, we utilized DNABERT model developed by Ji *et al*. [31] on human genomic data as an embedding layer to embed tokens into real-valued vectors. In the CNN model section, we employed a one-dimensional CNN to learn and classify the vectors outputted from the BERT embedding layer, thereby achieving the identification of m7G sites. The proposed GenoM7GNet surpasses existing state-of-the-art predictors in terms of accuracy, specificity, and Matthews Correlation Coefficient (MCC). The principal contributions of our work can be summarized as follows:

- A novel deep learning prediction model, GenoM7GNet, for rapid and accurate m7G sites identification.
- A new approach that integrates the BERT model based on nucleotide language as an embedding layer into m7G site prediction for the extraction of richer hidden information from RNA sequences.
- Without any extra information beyond RNA sequence, GenoM7GNet can accomplish high-performance m7G site prediction tasks.
- Compared to previous models, the method we propose demonstrates significant improvements across various evaluation metrics.

## 2 Materials and Methods

The proposed GenoM7GNet model comprises a series of processes including dataset preprocessing, embedding of pre-trained BERT model, training and classification through one-dimensional CNN, and model evaluation. The comprehensive workflow of the model is depicted in Figure 1. The detailed elaboration for each process is described in the following subsections.

**Fig. 1.**
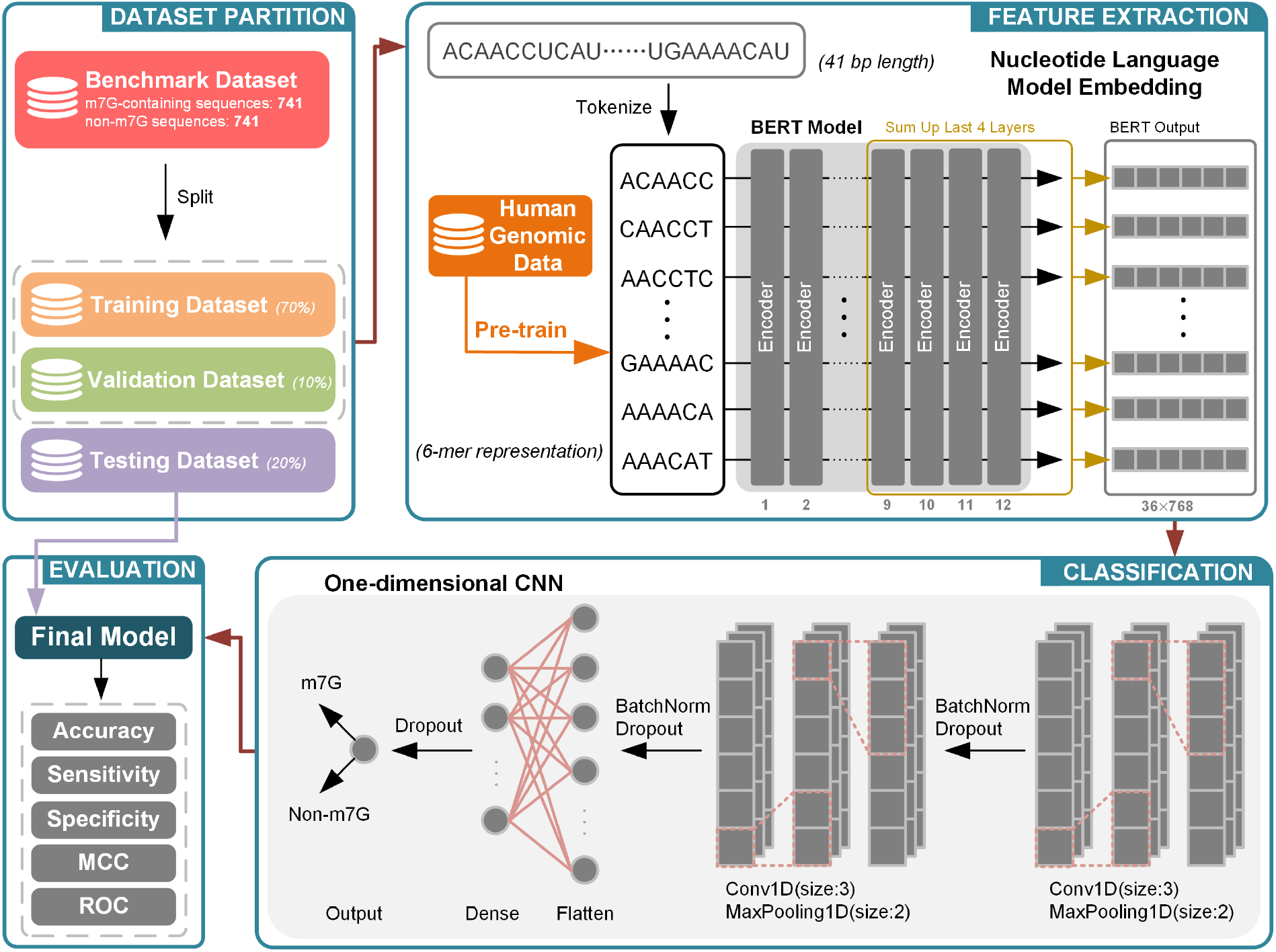
The flow chart of GenoM7GNet for predicting m7G sites. Firstly, we partitioned the benchmark dataset obtained from previous work into training, test, and validation sets, following a 70%, 20%, and 10% distribution, respectively. Subsequently, we utilize a BERT model pre-trained on human genome data as an embedding layer to extract information from RNA sequences. Then, we employ the 1D CNN architecture to learn the feature information derived from BERT and classify whether the sequences contained m7G sites. Finally, we use the test sets to evaluate the performance of the final model.

### 2.1 Dataset

In our work, we utilized the widely used dataset in previous research [20], [23], [24], [25], [26], [27]. This benchmark dataset is comprised of 1482 RNA sequences, each with a length of 41 nucleotides. It contains 741 positive samples (m7G sites) and an equal number of negative samples (non-m7G sites). The benchmark dataset D can be represented as

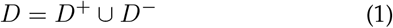

where *D*^+^ is the positive m7G site samples, and *D*^*−*^ the negative ones. *D*^+^ consist of sequences centered around m7G sites identified by Drummond *et al*. [32], originated from human HepG2 and HeLa cells. The negative samples contain non-m7G sites and are of equal length to the positive samples. To mitigate sequence homology bias, we employed the CD-HIT tool [33] to remove sequences with similarity greater than 80%. To intuitively observe the nucleotide base distribution between positive and negative samples, we employed an online Two Sample Logo tool [34] for analysis. The sequence logo shown in Figure 2 depicts the distribution of adenine (A), uracil (U), cytosine (C), and guanine (G) in the dataset.

**Fig. 2.**
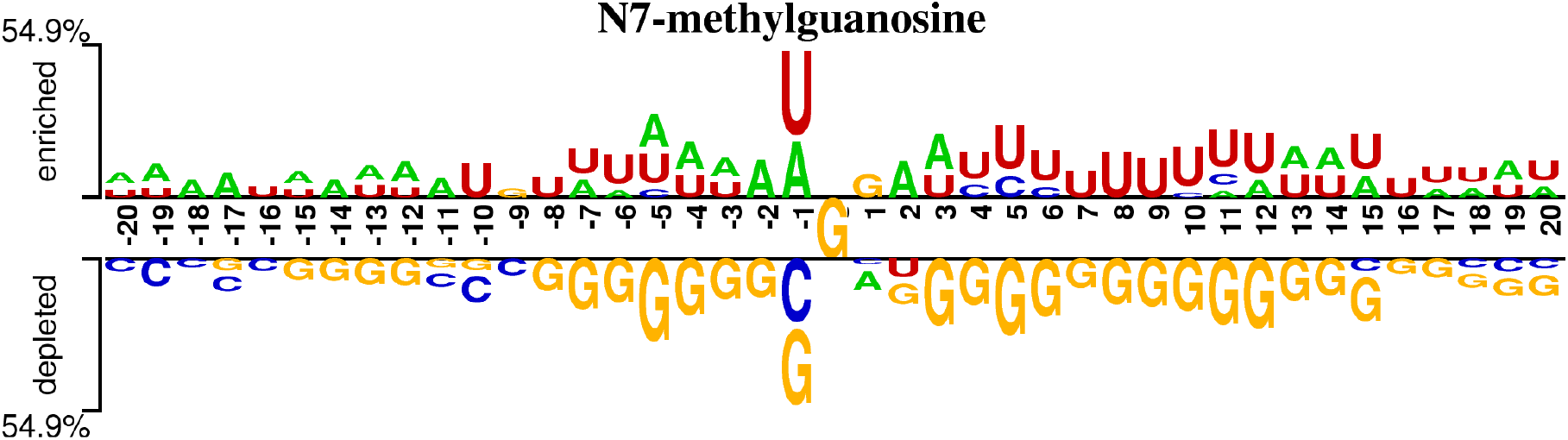
The distribution of nucleotides in the m7G site sample dataset. Proportional symbol sizes are used to illustrate the statistically significant differences between the two samples. Nucleotides are classified into two distinct categories: (i) those enriched in the positive samples and (ii) those that are depleted in the positive samples.

We first randomly partitioned the benchmark dataset, allocating 20% of the data as a testing set for final evaluation. To make optimal use of limited data and enhance the stability of the model, we employed an 8-fold crossvalidation approach for training. This choice is an empiric value from multiple experimental comparisons, where 8-fold cross-validation demonstrated markedly superior performance and generalizability compared to 5-fold or 10-fold cross-validation. Specifically, we evenly divided the remaining dataset into eight subsets, sequentially selecting one subset as the validation set and others for the training. The validation set is to evaluate the trained model with the goal of determining a final model that yields the optimal performance, while the test set is to assess this final model. The ratios among the training, validation, and testing sets within the benchmark dataset were 0.7:0.1:0.2, respectively.

### 2.2 Pre-trained BERT Model

#### 2.2.1 DNABERT

Biological sequences primarily consist of DNA, RNA, and protein sequences. They have been widely utilized in bioinformatics research and can be regarded as a form of language for conveying information between cells. Unlike natural languages, the information encoded in biological sequences is interacted at cellular level, making it complicated and difficult to understand. One of the primary challenges is to effectively extract and analyze information from these sequences. Word embedding is a method that maps words to real-valued vector representations, which can learn hidden features from large-scale corpora to convert words into suitable vector representations. For natural language processing (NLP), there are diverse widely used traditional word embedding methods, such as word2vec [35], [36], fastText [37], and GloVe [38]. Although these methods are efficient, they are inherently static and struggle to address the problem of polysemy. However, BERT utilizes the attention mechanism to pre-train unlabeled text bidirectionally. It is capable of generating dynamic word embeddings, where the same word or nucleotide is mapped to different real-valued vectors depending on its position in the sequence.

Inspired by the vector representation in nature language, we introduced DNABERT, which is an architecture based on BERT. DNABERT was pretrained on the human reference genome GRCh38.p13, with the training objective being the prediction of masked tokens using contextual cues. Numerous studies have demonstrated that features learned from DNA base sequences can be easily applied to RNA tasks [28], [39], [40], [41]. Compared to BERT models pre-trained on natural languages, DNABERT is superior to capture hidden information and features within nucleotide sequences.

#### 2.2.2 Tokenization

To enable us to handle with biological data, we utilized k-mer representation to tokenize nucleotide sequences as opposed to single bases. The k-mer representation is the concatenation of each nucleotide base with the subsequent continuous *k*-1 bases, forming a sequence of length *k*. For example, the 6-mer representation of the RNA sequence ‘ACAACCU’ is: {ACAACC, CAACCU}, while its 5-mer representation is: {ACAAC, CAACC, AACCU }. The k-mer representation, which provides richer contextual information for individual nucleotide bases, has been widely adopted to process biological sequences processing. In DNABERT, different k-mer representations (*k*=3-6) have been exploited to train four distinct models for downstream analysis. In our GenoM7GNet model, we opted to tokenize RNA sequences using 6-mer representation.

#### 2.2.3 BERT Embedding

After converting the sequences into 6-mer representation, a 41-length RNA sequence is segmented into 36 tokens. These tokens are then inputted into the pre-trained BERT model, where each token is transformed into a real-valued vector of dimension 768. Consequently, for each input RNA sequence, the BERT embedding layer converted it into a matrix of size 36×768. In our model, we adopted a strategy where parameters of the pre-trained BERT model remained fixed throughout the training phase. This means that we neither fine-tune nor train the model parameters individually, leaving all parameter learning to the downstream classifier. This decision was based on the fact that the pretrained BERT model already possesses strong nucleotide language information extraction capabilities, and due to the limitations in data volume, fine-tuning or participating in training offers limited improvements but more than doubles the time cost.

### 2.3 One-dimensional CNN Model

CNN is a deep learning model frequently employed for addressing computer vision problems, while also exhibiting exceptional performance in the field of bioinformatics [42], [43]. One-dimensional CNN, a variant of CNN, is particularly adept at handling sequence data and possesses robust abilities for capturing local features. In our research, we utilized the one-dimensional CNN model to perform feature learning on BERT embeddings and classify the presence of m7G sites based on the RNA sequences. We added a convolutional layer, a pooling layer, a fully connected layer to constitute the fundamental structure of our CNN models. To reduce the potential overfitting issues, we incorporate batch normalization and dropout strategies in the architecture.

In mathematical terms, the relationships among the convolutional layer and the Rectified Linear Unit (ReLU) layer are expressed as

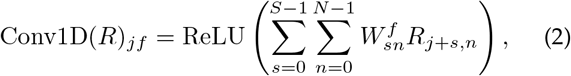

where *R* represents the input of one-dimensional CNN, *j* is the output position index, *f* is the convolutional kernel index, *s* denotes the kernel size and *n* is the number of channels. ReLU can be defined as

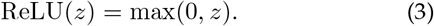

Batch normalization and dropout strategies can be employed to prevent the occurrence of vanishing gradients in the model. The mathematical definition of batch normalization is shown in Eq. (4).

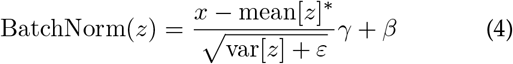

where mean[*z*] and var[*z*] represent the global mean and variance obtained during the training process, respectively. Following the two layers of 1D CNN is a fully connected neural network, the purpose of which is to flatten the CNN output and perform further feature extraction. Finally, we utlized the sigmoid function to scale the output layer value between 0 and 1, in order to predict whether a given sequence contains the m7G site. Equation (5) provides the mathematical expression of the sigmoid function.

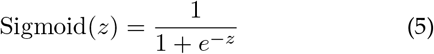

### 2.4 Performance Evaluation Metrics

In our research, we employed several commonly used metrics to assess the model’s predictive performance, including accuracy (ACC), sensitivity (SN), specificity (SP), Matthews Correlation Coefficient (MCC), and the receiver operating characteristic (ROC) curve.

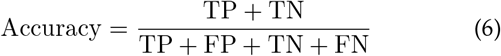

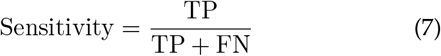

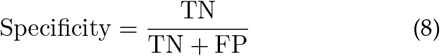

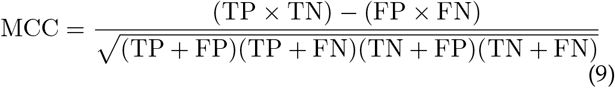

where TP, TN, FP, and FN represent the number of true positives (correctly identified positive samples), true negatives (correctly identified negative samples), false positives (negative samples misclassified as positive), and false negatives (positive samples misclassified as negative), respectively. Additionally, the ROC can reflect the predictor’s generalization capability to some extent. We employed AUC (Area Under the Curve) to represent the area beneath the ROC curve. The value closer to 1 indicates a higher level of robustness in the model.

### 2.5 Baseline Approaches

To evaluate the performance of our model, we compared GenoM7GNet against a range of existing baseline methods, including iRNA-m7G, m7GFinder, m7GPredictor, XG-m7G, m7G-IFL, m7G-DLSTM, THRONE, and m7G-autoBioSeqpy. iRNA-m7G, m7GFinder, and m7GPredictor employ SVM-based algorithms and leverage multiple feature subsets for the identification of m7G sites [20], [21], [22]. Both XG-m7G and m7G-IFL employ XGBoost-based approaches for the detection of m7G sites [23], [24]. m7G-DLSTM adopts deep learning techniques, utilizing Long Short-Term Memory (LSTM) sequences for feature extraction, learning, and classification [25]. THRONE is a machine learning-based three-layer ensemble predictor for m7G site identification [26]. m7G-autoBioSeqpy introduces autoBioSeqpy to employ a variety of deep learning methodologies for m7G site classification [27]. These baseline methods provide a comprehensive and multi-faceted benchmark for our evaluation.

## 3 Results

### 3.1 Hyperparameter Selection

Hyperparameter fine-tuning is crucial in deep learning, since it can optimize model performance. For the overarching parameters of our model, we chose a batch size of 64, a learning rate of 5e-6, and an epoch number of 100 as the training parameters for the GenoM7GNet model. To achieve better model performance, we adopted a linear learning rate warm-up strategy, where the learning rate linearly increases from null to the target value and then linearly decreases to zero throughout the training process. The training was conducted on a single NVIDIA A100 SXM4 40GB GPU, with the GenoM7GNet model taking approximately 30 minutes to train when the epoch count was set to 100. The model’s accuracy fluctuations in predicting the training and validation sets during the training process are depicted in Figure 3.

**Fig. 3.**
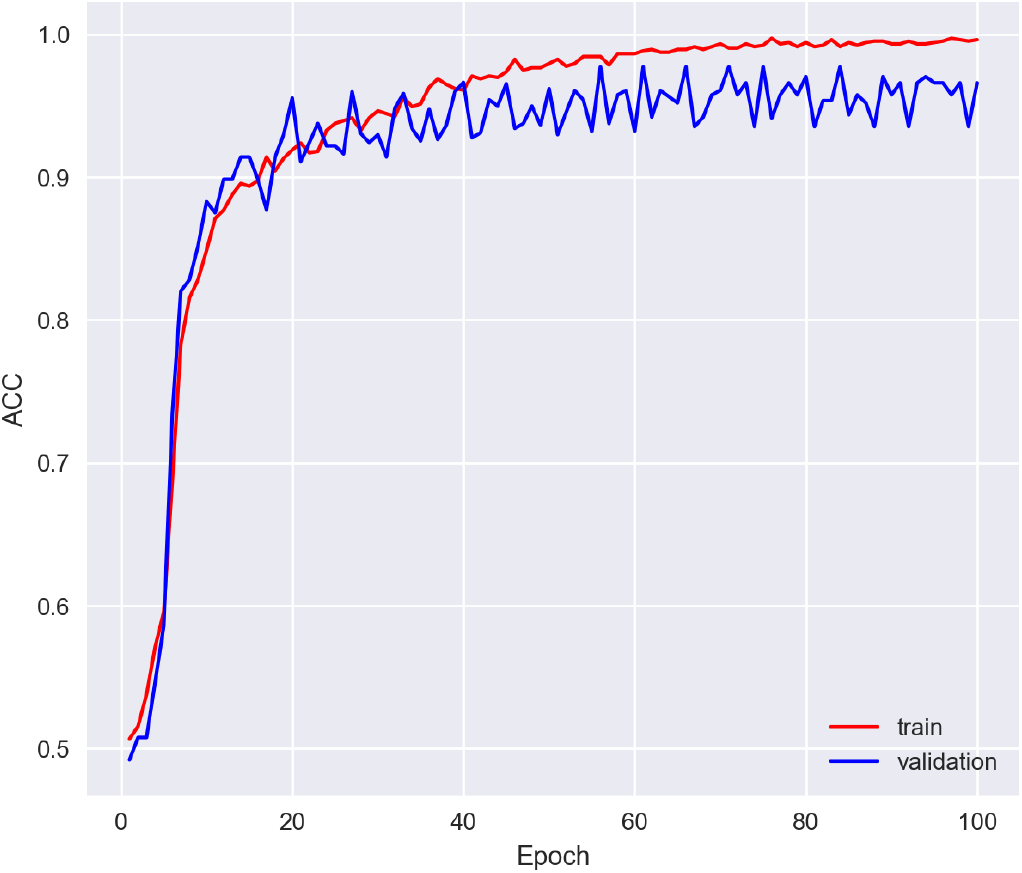
The learning curve of the GenoM7GNet model during the training process.

In our selection of hyperparameters for the one-dimensional CNN classifier, we assessed the model performance with distinct hyperparameters, including the number of convolutional layers, number of kernels, kernel size, maximum pooling size, and dropout rate post-convolution. Ultimately, we opted for the one-dimensional CNN with two convolutional layers, the first and second of which contain 72 and 200 kernels, respectively. Both layers employ a kernel size of 3, a maximum pooling size of 2, and a dropout rate of 0.2 post-convolution.

### 3.2 Model Performance Evaluation

Our GenoM7GNet model consists of a pretrained BERT model and a 1D CNN classifier, and it exhibited excellent performance in predicting m7G sites, with an accuracy of 0.953, sensitivity of 0.932, specificity of 0.976, and MCC of 0.907. In addition, the confusion matrix and ROC curve obtained from the test set evaluation of this model are shown in Figure 4, with an AUC value of 0.983. These results indicate that the BERT model, pretrained on the human genome data, possesses strong nucleotide language comprehension, enabling effective extraction of hidden features from biological sequences. Additionally, they also demonstrate the outstanding effectiveness and performance of the 1D CNN in local feature extraction and classification of sequence data.

**Fig. 4.**
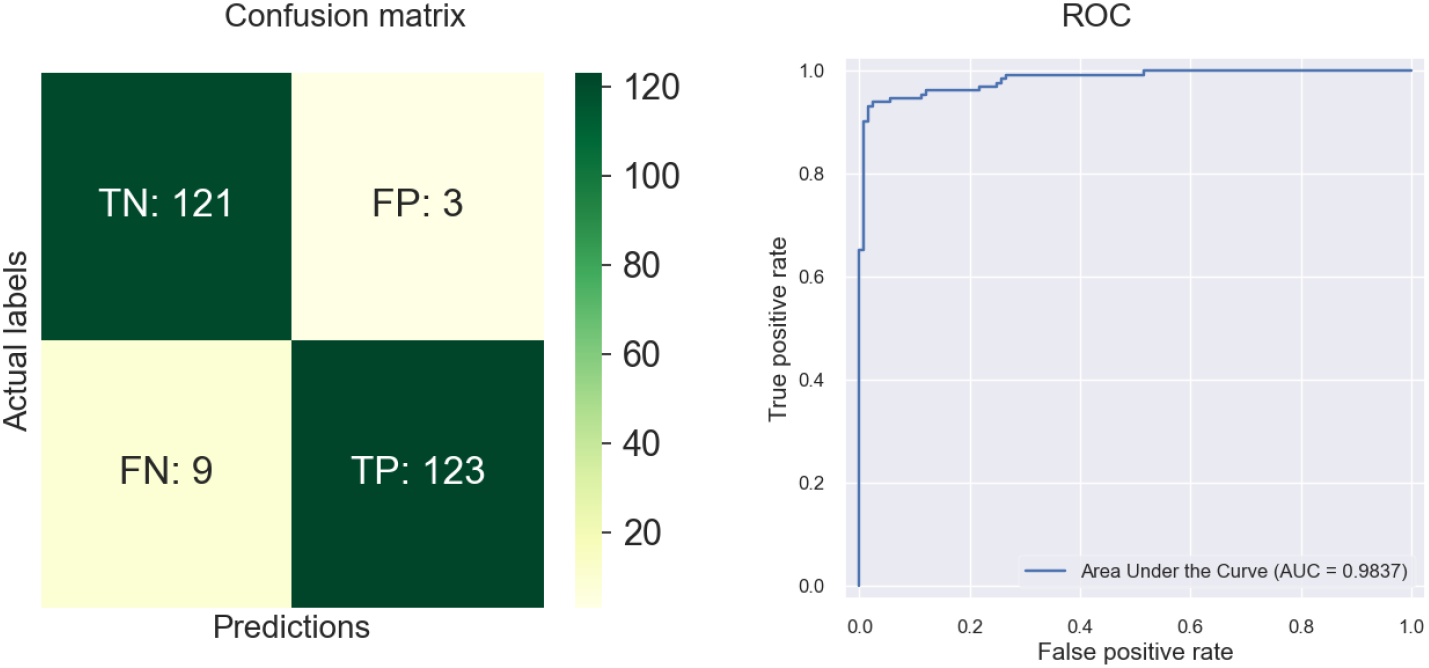
The confusion matrix and ROC curve graph of the GenoM7GNet model.

### 3.3 Visualization and Motif Discovery Based on BERT Embedding Layer

In our study, we utilized BERT as the embedding layer to extract hidden information from positive and negative samples. BERT is built on the Transformer architecture, with its core component being the attention mechanism. By visualizing the attention weights assigned to each token within the BERT embedding layer, we can gain an intuitive understanding of sample sequences related to m7G sites. Positions with higher attention scores may contain more critical information. We set the coordinate of the central ‘G’ base in the sequence to zero, which facilitates our observation of attention distribution surrounding the m7G site. Figure 5A and Figure 5B depicted the attention heatmaps for positive and negative samples over sequence positions, and Figure 5C presented the average attention score curve for both positive and negative samples. Although there are overarching similarities in the attention score for both samples, discrepancies are evident at specific positions.

**Fig. 5.**
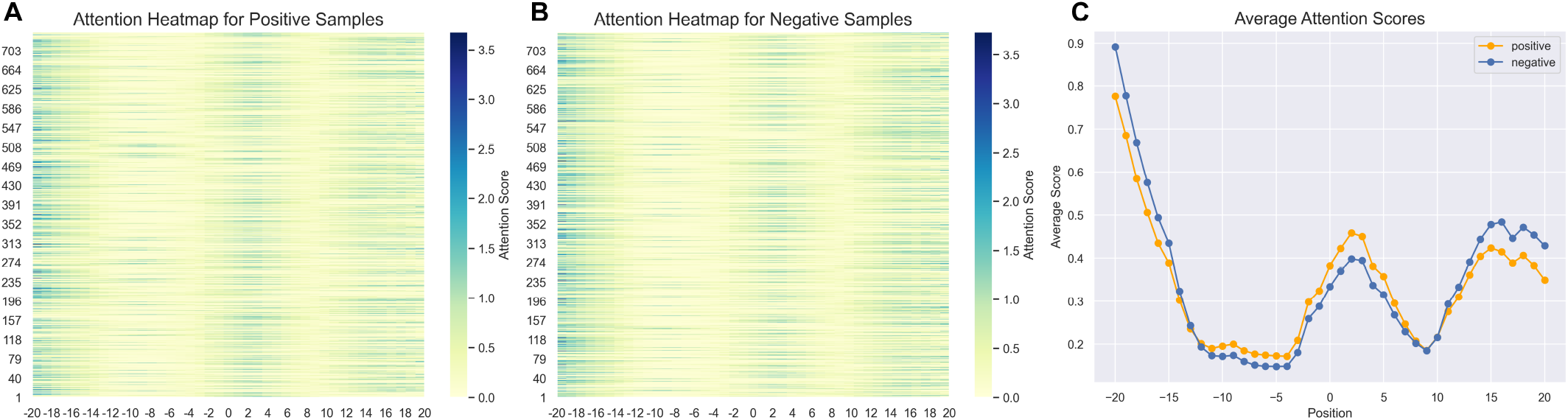
Visualization of attention scores at the m7G site for positive and negative samples. (A, B) Heatmaps depicting attention scores at various positions for both positive and negative samples. (C) Comparison of the average attention scores across different positions for positive and negative samples.

The attention scores can to some extent reflect the importance of sub-sequences. Hence, we can also leverage them to identify motifs of biological significance. Specifically, we first identified regions of high attention in the input sequence consecutively, and then employed the Aho-Corasick algorithm for multi-pattern matching to locate instances of motif patterns within the sequence. Subsequently, we conducted a hypergeometric test to find motifs significantly enriched in positive sequences, filtering out motif instances that have p-values less than 0.005. Finally, we aligned and merged similar motifs using the Smith-Waterman algorithm. Using a window size of 31, we identified three motif instances associated with m7G sites in RNA. These are represented as sequence logos in Figure 6.

**Fig. 6.**
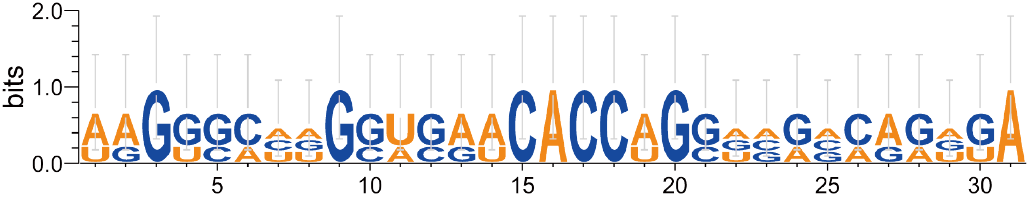
Sequence logo representation of the discovered motif at the m7G site.

### 3.4 Comparison to Previous Works on m7G Site Prediction

As previously mentioned, predicting m7G sites using RNA sequences is a valuable and appealing research topic, and experts have developed numerous advanced predictors. To demonstrate the capability of our model in predicting m7G sites, it is essential to compare with the previously most advanced predictors. In our study, we compared our GenoM7GNet model with several of the most advanced predictors available today, including iRNA-m7G, m7GFinder, m7GPredictor, XG-m7G, m7G-IFL, m7G-DLSTM, THRONE, and m7G-autoBioSeqpy [20], [21], [22], [23], [24], [25], [26],[27]. We applied the same datasets and evaluation metrics to guarantee an unbiased and fair comparison. The comparative results of the performance of each model are presented in Figure 7. Compared with the most advanced models previously available, our model’s accuracy, specificity, MCC, and AUC improved by 0.3%, 2.5%, 0.7%, and 0.9%, respectively. This significantly demonstrates the superior performance of our model in predicting m7G sites.

**Fig. 7.**
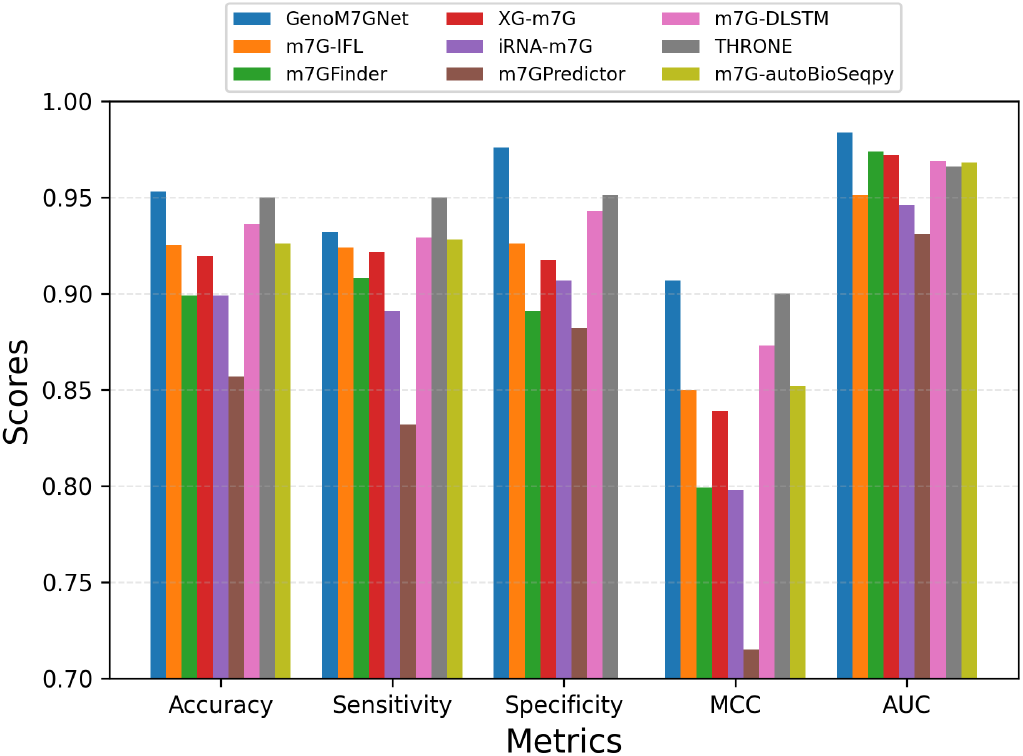
Performance comparison of our model with other state-of-the-art models.

To assess the stability of our model’s performance, we calculated 95% confidence intervals for accuracy, specificity, sensitivity, MCC, and AUC metrics, respectively, employing 30 data sets generated via cross-validation and random selection. Then, we used a t-distribution-based approach for these calculations, expressed by the formula

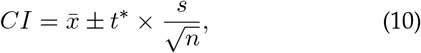

where 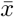 represents the sample mean, *s* the sample standard deviation, *n* the sample size, and *t*^*∗*^ the critical value from the t-distribution. To visually display the distribution and stability of the model performance metrics, we generated plots that include the 95% confidence intervals for each metric. As presented in Figure 8, a more intuitive understanding of the model evaluation results is depicted. By calculating the confidence intervals, we demonstrate the statistical significance of the model performance, which reveal clearly its potential variability range. However, it is important to note that these metrics are not entirely independent, which means that the chosen final model may not necessarily have all indicators within the confidence intervals.

**Fig. 8.**
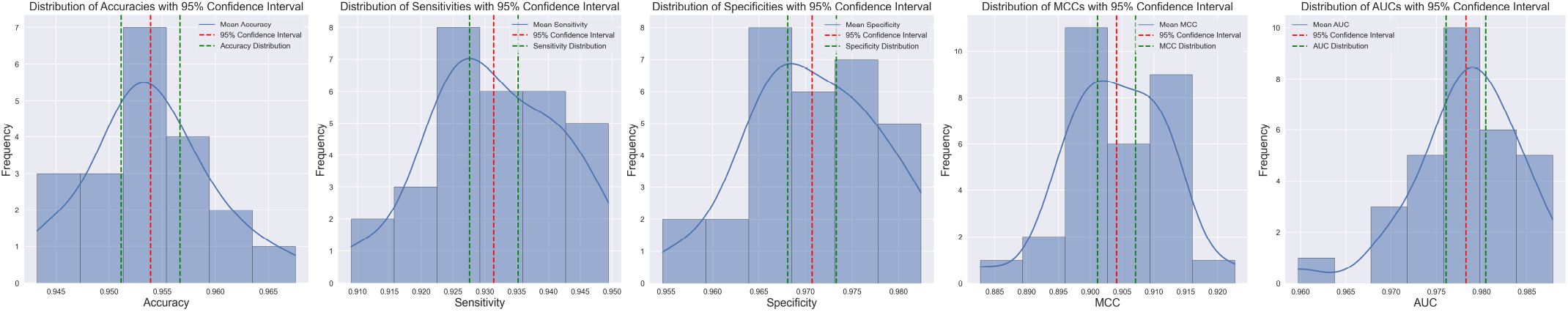
Distribution and 95% confidence interval of performance metrics for the GenoM7GNet model.

In addition, we tested our model on an independent dataset derived from m7GHub [21], which includes a more diverse and cross-species collection of m7G site data. We randomly selected a portion of m7G sites from different categories to form the positive samples. These positive samples consisted of 741 unique RNA sequences, each 41-bp in length, containing m7G sites. Negative samples were generated using the same method as the benchmark dataset, matching the positive samples in both quantity and length but without m7G sites. The test results demonstrated that our GenoM7GNet model achieved an accuracy, sensitivity, specificity, MCC, and AUC of 0.902, 0.880, 0.874, 0.880, and 0.964, respectively. We compared these results with those from available state-of-the-art models [20], [21], [22], [25], as shown in Figure 9. The findings indicate that our model consistently exhibits superior performance in terms of accuracy, specificity, MCC, and ROC curve. This suggests that GenoM7GNet not only has higher predictive performance but also strong generalization capabilities in predicting m7G sites.

**Fig. 9.**
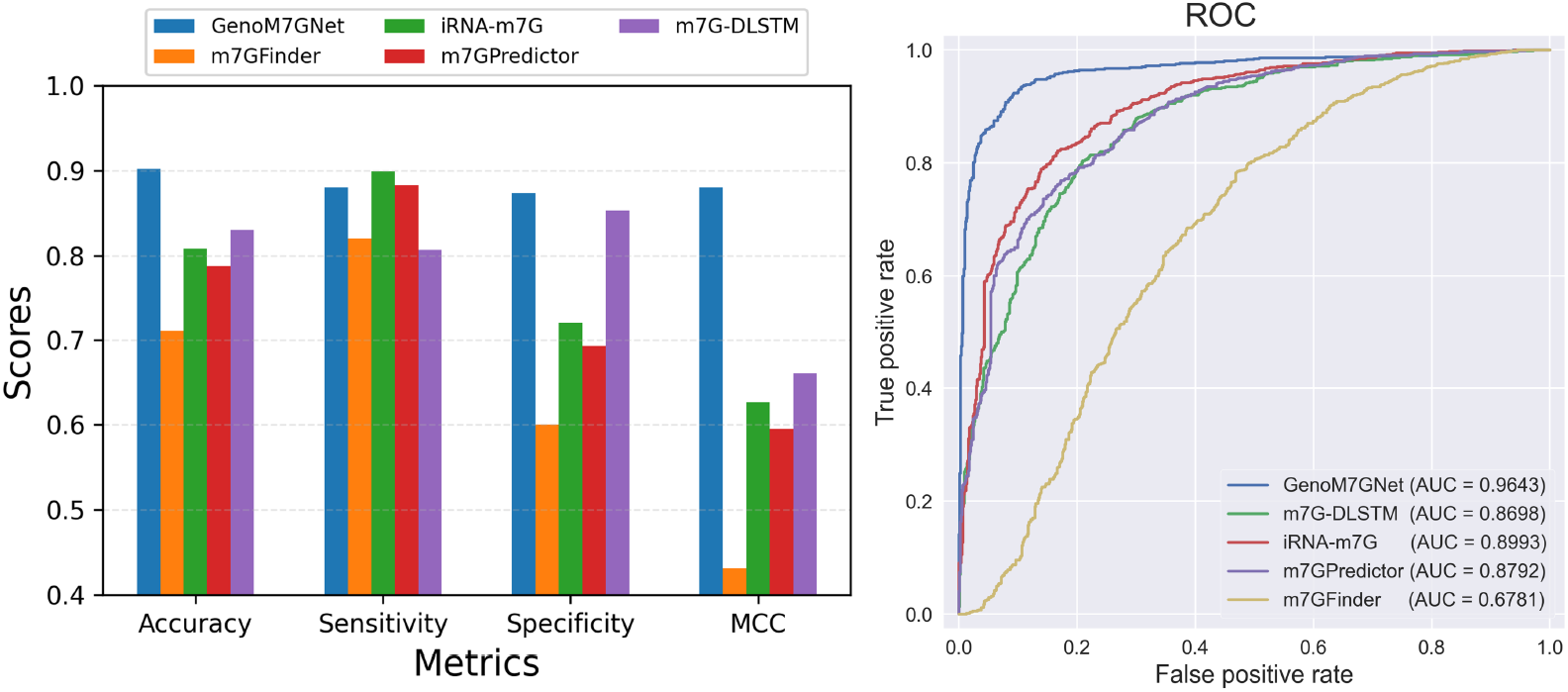
Performance comparison between our model and other available state-of-the-art models on the independent dataset.

### 3.5 Discussion on GenoM7GNet

In this sub-section, to better illustrate the function of each component of the GenoM7GNet model and the impact of hyperparameter selection, we conduct a discussion by deforming various parts of the model and adjusting the hyperparameters.

#### 3.5.1 Embedding Layer

To substantiate the enhanced capability of DNABERT pretrained on the human genome in extracting hidden information from RNA sequences, we conducted a comparative analysis between the performance of BERT pre-trained on natural language and DNABERT pretrained on different k-mers. It is clearly shown in Table 1 that DNABERT indeed exhibits a more powerful capacity to extract hidden information from RNA sequences. Additionally, the model pre-trained on 6-mer representations demonstrated superior performance.

**TABLE 1.** Comparison of pre-trained BERT models.

| Pre-trained BERT Model | ACC | SN | SP | MCC | AUC |
| --- | --- | --- | --- | --- | --- |
| BERT(natural languages) | 0.891 | 0.892 | 0.890 | 0.781 | 0.936 |
| DNABERT(3-mer) | 0.934 | **0.953** | 0.915 | 0.868 | 0.975 |
| DNABERT(4-mer) | 0.926 | 0.925 | 0.926 | 0.851 | 0.973 |
| DNABERT(5-mer) | 0.945 | 0.949 | 0.942 | 0.890 | 0.977 |
| DNABERT(6-mer) | **0.953** | 0.932 | **0.976** | **0.907** | **0.984** |

In our model, a BERT model pre-trained on the human genome is employed as the embedding layer. To evaluate its role and functionality, we compared it with two common embedding methods. The first is one-hot encoding, which maps different nucleotides into discrete vector representations. The second is the word2vec embedding model; we constructed a word2vec model trained on the same human genome data as DNABERT to be used as the embedding layer. This aimed to examine whether the dynamic BERT embedding exhibits excellent performance. The parameters of the word2vec model are detailed in Table 2.

**TABLE 2.**
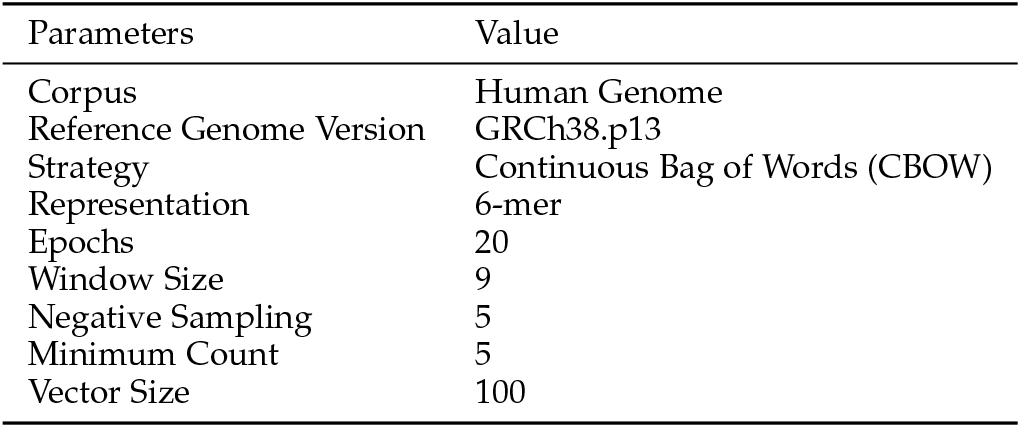
Training Parameters of Word2vec.

| Parameters | Value |
| --- | --- |
| Corpus | Human Genome |
| Reference Genome Version | GRCh38.p13 |
| Strategy | Continuous Bag of Words (CBOW) |
| Representation | 6-mer |
| Epochs | 20 |
| Window Size | 9 |
| Negative Sampling | 5 |
| Minimum Count | 5 |
| Vector Size | 100 |

We used various classifiers to learn and classify the outputs of these two embedding layers and evaluated them using the same metrics. The results are presented in Table 3. By comparing with the pre-trained BERT embedding in Table 1, it can be seen that models using BERT embeddings performed better overall compared to those using one-hot encoding and static word2vec embedding methods. This suggests that BERT embeddings dynamically generated from nucleotide sequences have superior capabilities to extract critical features to identify m7G sites. Moreover, it was observed that models using one-hot encoding generally outperformed those using word2vec embeddings. We speculate that this is due to the inability of word2vec to handle polysemous words and its ignorance of crucial features of nucleotide sequences during training.

**TABLE 3.**
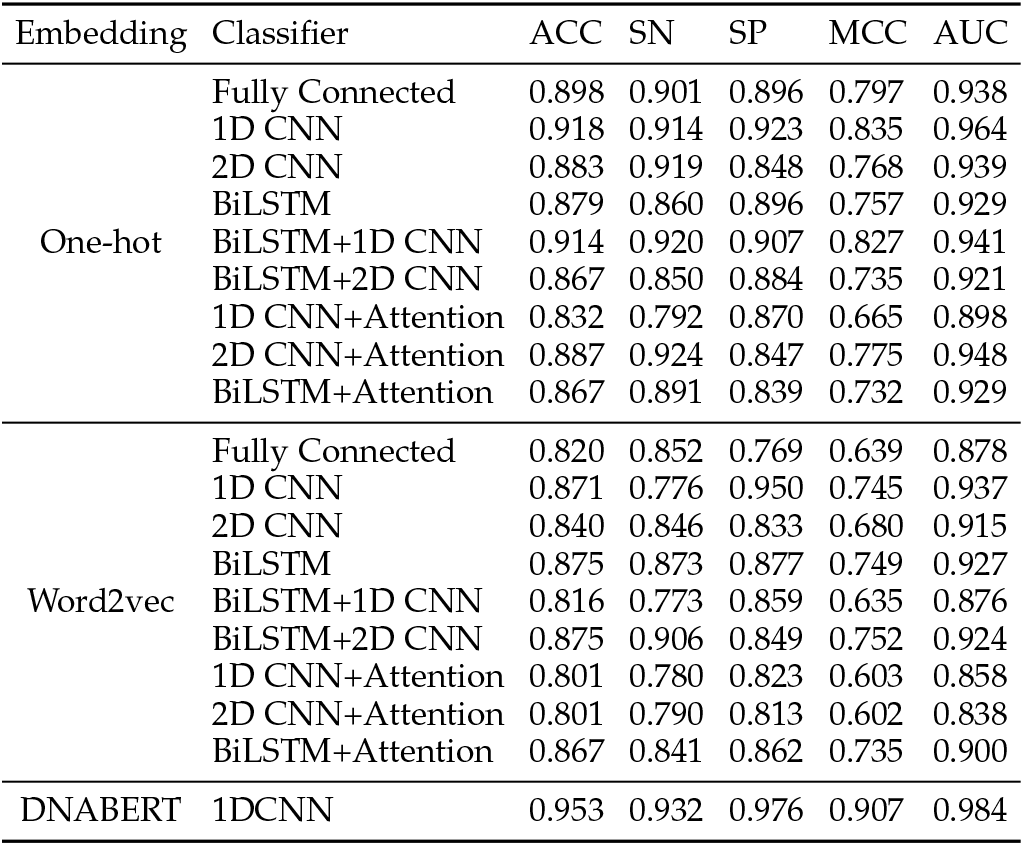
Performance comparison of models composed of different embedding methods and classifiers.

In GenoM7GNet, when generating the output vectors from the embedding layer, we chose to sum the last four layers of BERT and then output. The rationale behind this is that merging the outputs from multiple layers can yield a more comprehensive and rich representation. To verify its effectiveness, we constructed models by summing the output vectors from different numbers of final layers and subsequently tested the predictive accuracy. Table 4 shows the final tests results, proving that it exhibits optimal performance when the summing layer number is 4. This implies that its output can better integrate both local and global features of RNA sequences.

**TABLE 4.** Comparison of model performance with different summation layers in BERT embedding.

| Sum Layers | ACC | SN | SP | MCC | AUC |
| --- | --- | --- | --- | --- | --- |
| Last 1 layer | 0.902 | 0.900 | 0.904 | 0.805 | 0.961 |
| Last 2 layers | 0.926 | 0.912 | 0.939 | 0.852 | 0.962 |
| Last 3 layers | 0.941 | **0.969** | 0.911 | 0.884 | 0.969 |
| Last 4 layers | **0.953** | 0.932 | **0.976** | **0.907** | **0.984** |
| Last 5 layers | 0.926 | 0.906 | 0.945 | 0.852 | 0.968 |
| Last 6 layers | 0.910 | 0.905 | 0.915 | 0.820 | 0.942 |

#### 3.5.2 Classifier

To showcase the exceptional local feature learning and classification capabilities of 1D CNN, we attempted various advanced deep learning architectures for the identification of m7G sites, such as BiLSTM, CNN, attention mechanism, and their combinations. Through continuous hyperparameter adjustment and optimization of the various models and their blends, we found that the one-dimensional CNN demonstrated the most stable and outstanding performance, as shown in Table 5.

**TABLE 5.** Comparison of different classifiers.

| Classifier | ACC | SN | SP | MCC | AUC |
| --- | --- | --- | --- | --- | --- |
| Fully Connected | 0.863 | 0.845 | 0.850 | 0.724 | 0.928 |
| 1D CNN | **0.953** | 0.932 | **0.976** | **0.907** | **0.984** |
| 2D CNN | 0.910 | 0.895 | 0.924 | 0.820 | 0.946 |
| BiLSTM | 0.906 | **0.949** | 0.858 | 0.813 | 0.951 |
| BiLSTM+1D CNN | 0.902 | 0.865 | 0.938 | 0.806 | 0.930 |
| BiLSTM+2D CNN | 0.934 | 0.917 | 0.948 | 0.867 | 0.960 |
| 1D CNN+Attention | 0.910 | 0.908 | 0.913 | 0.820 | 0.953 |
| 2D CNN+Attention | 0.895 | 0.892 | 0.897 | 0.789 | 0.950 |
| BiLSTM+Attention | 0.906 | 0.893 | 0.920 | 0.813 | 0.929 |

For this outcome, our analysis suggests that the task may predominantly rely on local features within the sequence, making 1D CNNs more effective than models requiring longer-term dependencies (such as BiLSTM or attention mechanisms). Conversely, 2D CNNs, which are generally more suited for processing data with spatial relationships, might not effectively capture the dynamic features of time series. Additionally, the dataset in this task is not large, and simpler models may perform better than stacked models due to the latter’s tendency to overfit the training data. Simpler models are likely easier to generalize to unseen samples with limited data available.

To assess the effects of various parameters and strategies in the 1D CNN on model performance, we constructed prediction models with 1D CNN classifiers of different parameters. Firstly, we tested the impact of different numbers of layers and sizes of convolutional kernels on model predictive accuracy. The results of which are shown in Figure 10. It was observed that the models exhibit optimal performance when the number of layers is set to either two or three. This phenomenon can be attributed to the fact that different convolutional layer depths extract and analyze features at varying levels of abstraction. Yet an excessive number of layers may complicate the model, leading to overfitting and information loss. Moreover, 1D CNNs with a kernel size of 3 generally outperformed with other sizes. This is presumably because, in this task, a kernel size of 3 is sufficiently capable of capturing local contextual features while avoiding the introduction of excessive irrelevant information, thus preventing interference and overfitting.

**Fig. 10.**
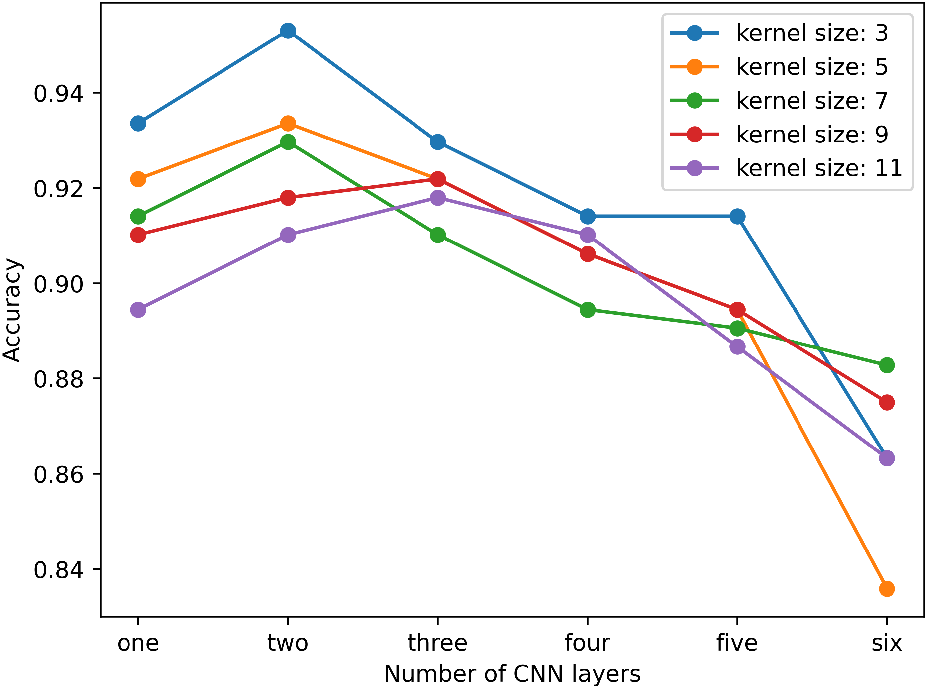
Line chart of model accuracy corresponding to different numbers of CNN layers and convolution kernel sizes.

We also compared the effects on model performance of different dropout rates in the CNN with or without normalization. As can be seen from Figure 11, incorporating dropout strategies can effectively prevent overfitting, thereby improving the model’s generalization performance. However, a high dropout rate may prevent the model from obtaining sufficient information. Batch normalization can effectively address the issue of internal covariate shift in the model, thereby improving model performance and accelerating model training speed.

**Fig. 11.**
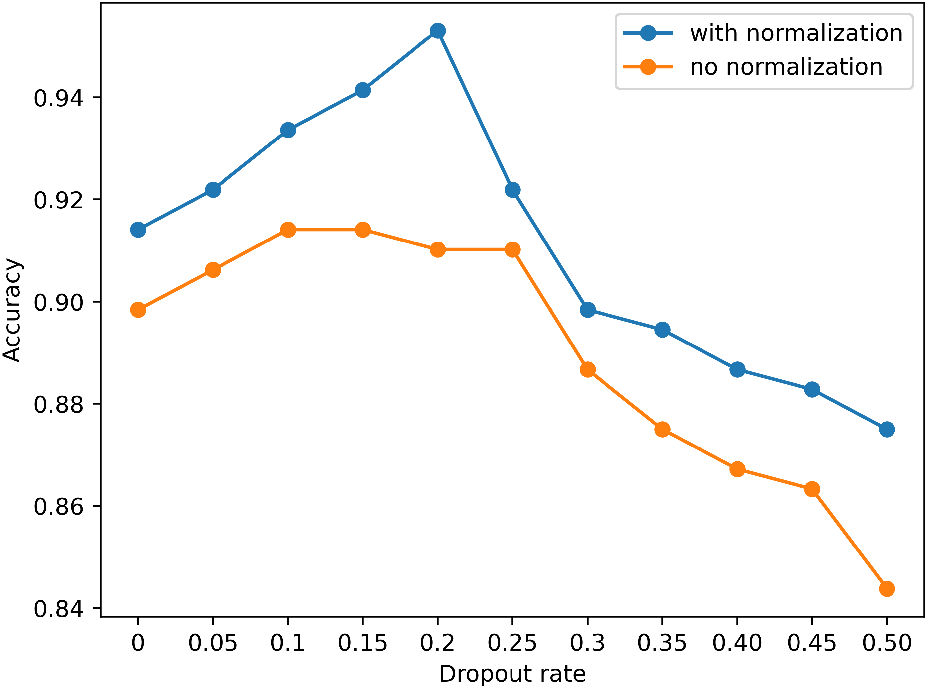
Line graph depicting the impact of dropout rates and batch normalization on model accuracy in the 1D CNN classifier.

## 4 Conclusion

To efficiently identify the m7G sites, this paper proposes a novel deep learning model called GenoM7GNet. By utilizing a BERT model pre-trained with the nucleotide language, more rich and reliable hidden information and features were extracted. Compared to BERT models pre-trained on natural language, our model demonstrates superior capability in extracting RNA sequence information, making it more suitable for biological sequence processing and analysis. Predictive results indicated that our proposed GenoM7GNet model exhibits efficient and outstanding performance in the identification of m7G sites, surpassing other state-of-the-art models. This model will assist medical professionals in drug development and cancer treatment, advancing the further development of precision medicine.

However, our approach has certain limitations. Due to the limited current research on the m7G site, the dataset we can utilize is relatively small. Consequently, our model might not be able to gather more biological insights on the m7G sites from the results. In future research, we plan to gather more extensive and higher-quality m7G site data, and adjust our model training strategies, while incorporating more advanced deep learning methods to further enhance the accuracy and efficiency of prediction.

---

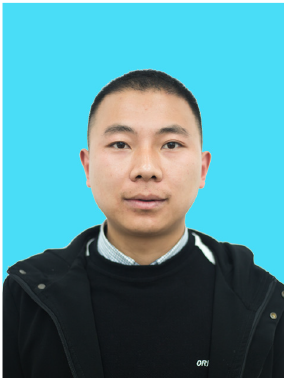

**Chuang Li** (Member, IEEE) received the Ph.D. degree in computer science from Hunan University, China. He is currently an associate professor of Hunan University of Technology and Business. He studied at Biomedical Informatics Lab at NTU from 2017 to 2018. His major research areas include high-performance computing, edge computing, bioinformatics. He has published research articles in Bioinformatics, BMC bioinformatics, IEEE Transactions on Services Computing.

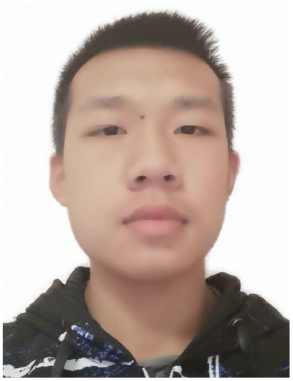

**Heshi Wang** received his bachelor’s degree from Central South University of Forestry and Technology, Changsha, China, in 2022. He is currently pursuing a master’s degree in the School of Computer Science at Hunan University of Technology and Business, Changsha, China. His research interests include bioinformatics, deep learning, and machine learning.

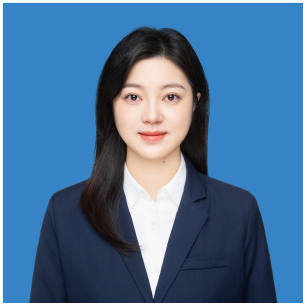

**Yanhua Wen** received the PhD degree in electronic from the Polytech’ Nantes, France, in 2013. She was a member of the Institut d’É lectronique et de TÉlÉcommunications de Rennes (IETR), one of the State Key Laboratory of France. She is currently a lecturer in Computer Science at Hunan University of Technology and Business. Her major research interests are image reconstruction, signal processing, electro-magnetic and high-performance computing. She has published many high-level academic papers in journals and conferences such as PIER, Journal of physics, Antennas and Propagation and USNC-URSI National Radio Science Meeting and so on.

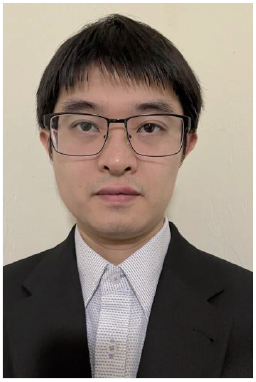

**Rui Yin** is currently an assistant professor at the Department of Health Outcomes and Biomedical Informatics, University of Florida. He completed his postdoctoral training at the Harvard Medical School. Before that, he received his Ph.D. degree from the School of Computer Science and Engineering, Nanyang Technological University. His research interests mainly focus on AI-driven precision medicine to improve public health outcomes and equity, where he develops novel machine learning approaches to address biomedical problems, such as in RNA viruses and infectious diseases, rare disease and cancer study, and health equity. He has published over 25 papers and served as an editorial committee member for several journals and conferences.

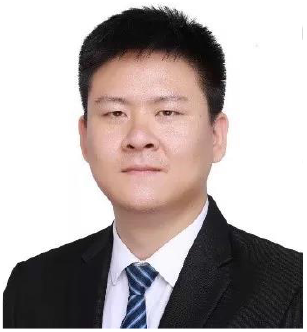

**Xiangxiang Zeng** (Senior Member, IEEE) received the BS degree in automation from Hunan University, China, in 2005, and the PhD degree in system engineering from the Huazhong University of Science and Technology, China, in 2011. He is an Yuelu distinguished professor with the College of Information Science and Engineering, Hunan University, Changsha, China. Before joining Hunan University in 2019, he was with the Department of Computer Science in Xiamen University. His main research interests include membrane computing, neural computing and bioinformatics.

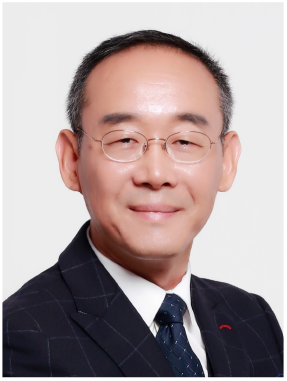

**Keqin Li** (Fellow, IEEE) is a SUNY Distinguished Professor of computer science with the State University of New York. He is also a National Distinguished Professor with Hunan University, China. His current research interests include cloud computing, fog computing and mobile edge computing, energy-efficient computing and communication, embedded systems and cyber-physical systems, heterogeneous computing systems, big data computing, high-performance computing, CPU-GPU hybrid and cooperative computing, computer architectures and systems, computer networking, machine learning, intelligent and soft computing. He has authored or coauthored more than 780 journal articles, book chapters, and refereed conference papers, and has received several best paper awards. He holds over 60 patents announced or authorized by the Chinese National Intellectual Property Administration. He is among the world’s top 10 most influential scientists in distributed computing based on a composite indicator of Scopus citation database. He has chaired many international conferences. He is currently an associate editor of the ACM Computing Surveys and the CCF Transactions on High Performance Computing. He has served on the editorial boards of the IEEE Transactions on Parallel and Distributed Systems, the IEEE Transactions on Computers, the IEEE Transactions on Cloud Computing, the IEEE Transactions on Services Computing, and the IEEE Transactions on Sustainable Computing.

